# TAG.ME: Taxonomic Assignment of Genetic Markers for Ecology

**DOI:** 10.1101/263293

**Authors:** Douglas Eduardo Valente Pires, Francislon Silva Oliveira, Felipe Borim Correa, Daniel Kumazawa Morais, Gabriel Rocha Fernandes

## Abstract

1.

**Background:** Sequencing of amplified genetic markers, such as the 16S rRNA gene, have been extensively used to characterize microbial community composition. Recent studies suggested that Amplicon Sequences Variants (ASV) should replace the Operational Taxonomic Units (OTU), given the arbitrary definition of sequence identity thresholds used to define units. Alignment-free methods are an interesting alternative for the taxonomic classification of the ASVs, preventing the introduction of biases from sequence identity thresholds.

**Results:** Here we present TAG.ME, a novel alignment-independent and amplicon-specific method for taxonomic assignment based on genetic markers. TAG.ME uses a multilevel supervised learning approach to create predictive models based on user-defined genetic marker genes. The predictive method can assign taxonomy to sequenced amplicons efficiently and effectively. We applied our method to assess gut and soil sample classification, and it outperformed alternative approaches, identifying a substantially larger proportion of species. Benchmark tests performed using the RDP database, and Mock communities reinforced the precise classification into deep taxonomic levels.

**Conclusion:** TAG.ME presents a new approach to assign taxonomy to amplicon sequences accurately. Our classification model, trained with amplicon specific sequences, can address resolution issues not solved by other methods and approaches that use the whole 16S rRNA gene sequence. TAG.ME is implemented as an R package and is freely available at http://gabrielrfernandes.github.io/tagme/

## 3. Introduction

The study of microbial community structure and variability has revolutionized our understanding of human health and disease, the discovery of novel species and their role in environments across the globe [1, 2].

Sequencing of amplified genetic markers, including the 16S ribosomal RNA gene, has been extensively used to characterize microbial community composition [3]. The cost-effectiveness of amplicon-based microbiome studies may explain its popularity. The existence of specialized databases in the taxonomic information of marker genes - such as Ribosomal Database Project [4], Greengenes [5], SILVA [6] - also reinforces the relevance of such an approach.

The organization of sequences into Operational Taxonomic Units (OTUs) (*i.e*., group of sequences that correspond to a given level of dissimilarity) traditionally precedes the taxonomic classification [7]. However, the OTU approach ignores specific variation by summarizing them into arbitrary identity thresholds [8]. The DADA2 [9] and Deblur [10] are approaches that consider the Amplicon Sequence Variants (ASVs) as a representation of each organism in the sequenced microbiome. In this context, alignment-free methods are an interesting alternative for the taxonomic classification of the ASVs, preventing the introduction of biases from the sequence identity thresholds. Machine learning-based methods, such as RDP Classifier[11], are widely used for the taxonomic assignment. For instance, DADA2 uses a predictive model trained with the Naïve Bayes algorithm to classify sequences up to the genus level, however the species identification relies on the exact sequence match against a reference database, which makes the procedure slow and insensible to fine-scale variations. Supervised learning techniques can leverage the wealth of genetic marker sequencing information available on the literature, being capable of generalization (predicting unseen data) as well as providing scalable approaches. In this work, we propose TAG.ME, a scalable alignment-independent and amplicon-specific method for taxonomic assignment based on genetic markers.

## 4. Implementation

### Database preparation

TAG.ME is fully customizable, allowing users to tailor personalized predictive models. A package of scripts and a tutorial are available at http://gabrielrfernandes.github.io/tagme/. The downloadable model files were developed using the SILVA database, version 132. Figure S1 of Supplementary Materials depict the complete TAG.ME pipeline. The user can build their training databases using a FASTA file containing the marker genes sequences of interest and a taxonomic description per sequence.

After database creation, amplicons are computationally generated based on user-selected primers, followed by a dereplication [12] step to obtain a non-redundant set of sequences. The last common ancestor (LCA) of each group of redundant sequences is calculated. K-mer frequencies are computed and used as features for training the classification models. k-mers of 3, 4 and 5 nucleotides were tested to identify the best k value to construct the models. Shorter k-mers (k < 4) generated less discriminatory models because of the fewer number of features; while 5-mer model relies on sparse matrices that do not fit well into the Random Forest algorithm. K=4 was the most discriminatory word length and was therefore used in this work. A k-mer frequency table is calculated and used during supervised learning.

### TAG.ME pipeline - Supervised Learning

TAG.ME uses the random forest ensemble algorithm to train its amplicon-specific predictive models. Novel models can be trained upon request via the website https://gabrielrfernandes.github.io/tagme, or locally via the package of scripts available for download. For each taxonomic level, a predictive model is trained, where each taxon represents a class. During training of each taxonomic level, 40% of classes are randomly filtered out. This composes a true negative set. The remaining set is further divided in training (60% of the remaining) and testing (40% of the remaining), making sure all classes have representatives in each of the sets. This procedure is repeated 20 times, and average performance metrics are derived to calculate sensitivity and specificity metrics.

### Prediction assignment and sensitivity/specificity tradeoffs

As a multi-level classification system, TAG.ME employs a bottom-up procedure for taxonomy assignment as follows. Starting from the deepest level (Species), it assesses the probability of assignment to each class by the random forest classifier. The classification score is calculated based on the two top-ranked classes according to the formula: Score = log2(P(α)/P(β))×P(α), Where, P(α) is the probability for the top-ranked class prediction, α; and P(β) is the probability for the second-ranked class, β. The chosen value for the score influences sensitivity and specificity tradeoffs (Supplementary Table 1), which can be set by the user. By default, TAG.ME taxonomic assignment is set to deliver a specificity of at least 80%. If the Score is higher than the threshold, a taxon is assigned. Otherwise, TAG.ME raises the taxonomy to the next taxonomic level and repeats the process.

### Models Optimization

The number of features and class representativity used to build the models are also optimized in order to reduce time and memory requirements during prediction and guaranteeing scalability. The relative importance (Mean Decrease Gini) was calculated, and ordered, for each of the 136 possible 4-mers during model construction. To select the number of features to be used to train the final model, we summed the relative importance of each feature until it reaches 70%, 80%, and 90%. The Area Under the Receiver Operator Characteristic Curve (AUROC) was calculated for each condition and compared. The model containing fewer features is selected unless there is an improvement in the AUROC (p<0.05). The same procedure was used to determine the maximum number of points to describe one class. We tested models limiting the number of points to the 85, 90, 95 and 100 percentile of the classes sizes. The AUROC was calculated and compared as previously described.

### Benchmark

The rRNA 16S genes unaligned FASTA sequences and taxonomic lineage for each entry were downloaded from the Ribosomal Database Project website (release 11.5) [4]. The 3,196,041 sequences were filtered to remove entries without the full lineage description, keeping only 276,214 entries with information up to species level. The dataset was used as the reference for the amplicon generation, dereplication, and LCA identification, as described in Database preparation. The 42,069 non-redundant amplicon sequences were used as reference dataset for benchmarking and submitted to the taxonomic classification using popular pipelines with their default parameters: 1) Utax (USEARCH v10.0.240) [13]; 2) QIIME (version 1.9.1) [7] using UCLUST [13] method with two different databases - Greengenes 13.5 [5] and SILVA 129 [6] -, and RDP[11] as method; 3) the standalone RDP Classifier (version 2.2) [11]; 4) QIIME2 feature-classifier [14] using the Naïve Bayes algorithm to train the same database used in TAG.ME models; and 5) DADA2 (version 1.9) [9]. The unclassified or inexact classifications were excluded from the reference dataset before the comparison. The correct assignments are those which the prediction exactly matches the reference classification, and the misassignments are those that do not match. The unclassified sequences represent entries that have a precise taxonomic description in the reference set, but could not be identified by the classification tool. The benchmark dataset was used to evaluate the running time of the tools as well.

Another benchmark was performed using the Mockrobiota [15] dataset. The expected sequences present in 6 sets - Mock-3 [16], Mock-12 [9], Mock-13 [17], Mock-16 [18], Mock-18 [19] and Mock-20 [20] - were used as reference for an *in silico* extraction of the amplicon equivalent to the tested model: 515F-806R. Each unique sequence was classified using TAG.ME, DADA2, and QIIME2 feature-classifier using the Naïve Bayes method for comparison.

The overclassification rate - or false positive fraction - was compared to the measures obtained from the publication that suggests the parameters to optimize the taxonomic classification using QIIME2 [14]. The "novel taxa” data was used to train and test the TAG.ME models. The experiment consists in predicting taxonomy to a class that is not present in the training set - working as a Negative dataset.

### Test data sets

DADA2 classification into species-level relies on exact sequence match against the database. The benchmark dataset was obtained from RDP, which is the same database used for DADA2 Species assignment, leading to better accuracy than TAG.ME in that taxon. Two published datasets were used to evaluate the performance of sequenced data. The dataset available under the accession PRJEB19103 [21] represents the human gut environment, and accession PRJEB15671 [22] represents the root ecosystem. The amplicon sequence variants (ASVs) were identified using the DADA2 workflow. Each ASV was classified by DADA2 using the training set from RDP and SILVA. TAG.ME used the 515F-806R model (V4 region) to perform the taxonomic assignment.

### Code availability

TAG.ME is freely available as an R package via GitHub (https://gabrielrfernandes.github.io/tagme). Tutorials, as well as a help guide, were developed and are available on a user-friendly web interface. TAG.ME is fully integrated with other well established pipelines, such DADA2’s and Phyloseq [23].

## 5. Results and Discussion

### Model performance

This Results section will focus on the experiments using the model 515F-806R (V4 region of the 16S rRNA gene), however the results are reproducible and comparable to other amplicon models as shown in Supplementary Figure 2. The definition of the number of features and class coverage was determined based on the improvement of the AUROC across the different tests. The Supplementary Figure 3 shows the distribution of AUROC in the different conditions: 1) 70%, 80% and 90% cumulative importance of variables, and 2) relative class representativity of 0.85, 0.90, 0.95, and 1. The distribution of importance per feature shows that some elements have a higher influence in distinguishing between taxa, as shown in Figure 1A. This observation supports the limitation of the number of features to train the models. In some cases, such as in the species taxon, increasing the number of features degrades performance, evidencing the incorporation of noise by low importance features (Supplementary Figure 3A). After defining an optimized feature set, the 40/60-split procedure described in the “Supervised Learning” subsection was used to calculate the sensitivity and specificity of the final model. The Receiver Operator Characteristic (ROC) curves pictured in Figure 1B displays the high efficacy of the TAG.ME in predicting taxonomy in every level.

**Figure 1.**
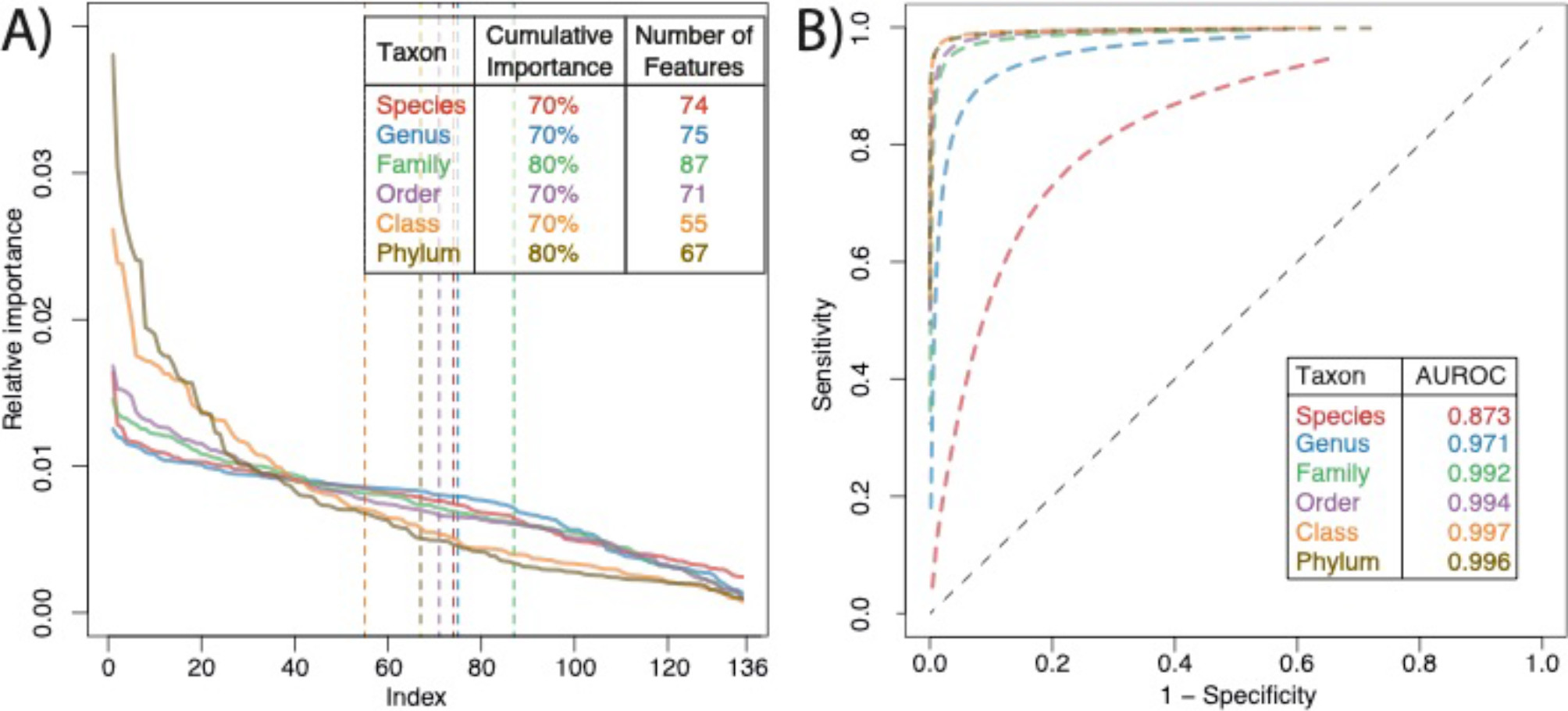
(A) The relative importance of each 4-mer was summed to define the features used to build the prediction model. The vertical traced lines determine the last selected feature according to the their cumulative importance. (B) The ROC curves for each Taxon demonstrates the high accuracy of all TAG.ME models.

The performance measures were compared to the methods and parameters suggested by QIIME2 feature-classifier [14]. The F-measure, precision, recall and overclassification Rate were calculated for each taxonomic rank - for TAG.ME models - and obtained from the classification evaluation performed, suggested and published by Bokulich et. al. The mentioned manuscript indicates methods and parameters to optimize QIIME2 classification, four of the most balanced results are compared in Table 1. TAG.ME showed the overclassification rate limited to the 20% which is expected as the specificity is set to 0.8; while machine learning QIIME2 optimized methods - Naïve Bayes and RDP - deliver a higher rate of overclassification. The UCLUST algorithm showed a lower overclassification rate - 6% -, on the other hand, had the highest underclassification rate. All the performance measures for TAG.ME are reported in Supplementary Table 2.

**Table 1.**
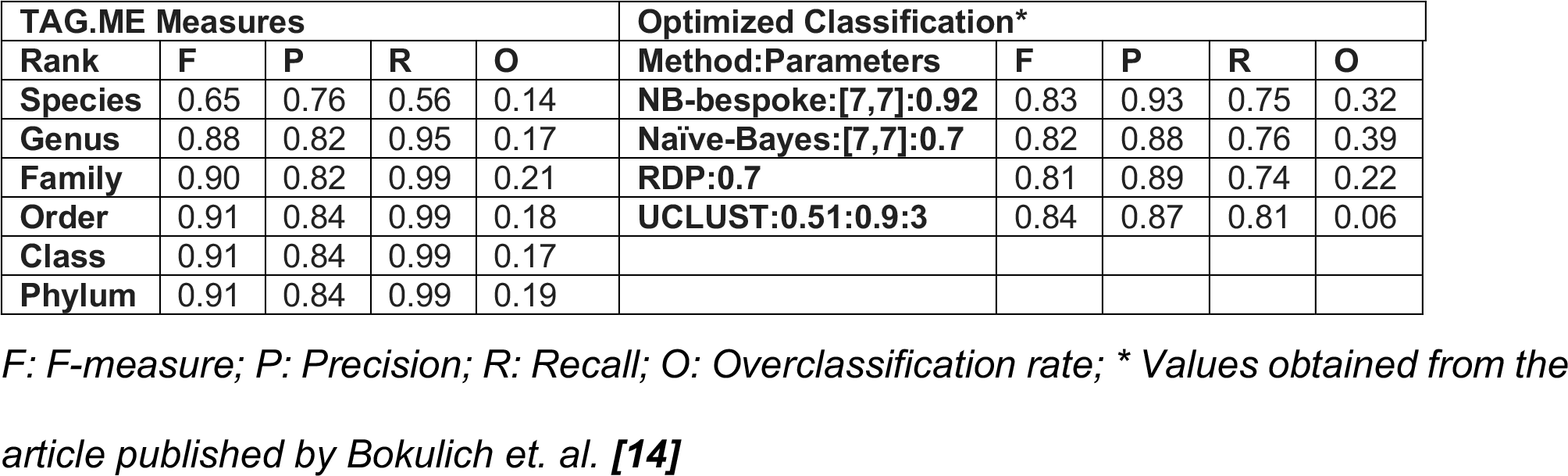
TAG.ME performance measures compared to other methods.

| TAG.ME Measures |  |  |  |  | Optimized Classification* |  |  |  |  |
| --- | --- | --- | --- | --- | --- | --- | --- | --- | --- |
| Rank | F | P | R | O | Method:Parameters | F | P | R | O |
| Species | 0.65 | 0.76 | 0.56 | 0.14 | NB-bespoke:[7,7]:0.92 | 0.83 | 0.93 | 0.75 | 0.32 |
| Genus | 0.88 | 0.82 | 0.95 | 0.17 | Naïve-Bayes:[7,7]:0.7 | 0.82 | 0.88 | 0.76 | 0.39 |
| Family | 0.90 | 0.82 | 0.99 | 0.21 | RDP:0.7 | 0.81 | 0.89 | 0.74 | 0.22 |
| Order | 0.91 | 0.84 | 0.99 | 0.18 | UCLUST:0.51:0.9:3 | 0.84 | 0.87 | 0.81 | 0.06 |
| Class | 0.91 | 0.84 | 0.99 | 0.17 |  |  |  |  |  |
| Phylum | 0.91 | 0.84 | 0.99 | 0.19 |  |  |  |  |  |

*F: F-measure; P: Precision; R: Recall; O: Overclassification rate; \* Values obtained from the article published by Bokulich et. al. [14]*

Additionally, models were constructed to evaluate the overclassification rate using the same training and testing dataset as used to represent “novel taxa” in Bokulich’s work. TAG.ME had good performance when compared to other methods, especially at phylum and class levels as showed in Supplementary Figure 4. Although Table 1 reports different values and better overall measures, this is explained by the number of points representing each class used to train the model. As shown in Supplementary Figure 3A, the model performance is impacted by the class representativity. In most of the scenarios, as more points are representing one taxon in the construction of the trees, better is the performance.

### Comparison with other tools

To better assess TAG.ME’s performance, we compared our approach with five well-established methods - 1) UTAX (From USEARCH package) [13], 2) RDP Naïve Bayes classifier [11], 3) QIIME [7] (using Greengenes [5] and SILVA [6] databases as reference, and RDP Classifier [11] as algorithm), 4) QIIME2 feature-classifier [14] and 5) DADA2 [9] (using SILVA [6] and RDP [4] as databases) - on a large scale test, using the RDP sequences prepared to serve as benchmark as assignment target. Figure 2 depicts the performance of each method on assigning the different taxonomic levels of the benchmark dataset from domain to species. We show TAG.ME performed as well as or better than alternative approaches for all levels. It is noteworthy that DADA2’s performance on species level, despite being slightly superior, occurred because the method finds exact matches in the reference database: the RDP dataset, the same used to construct the benchmark dataset.

**Figure 2.**
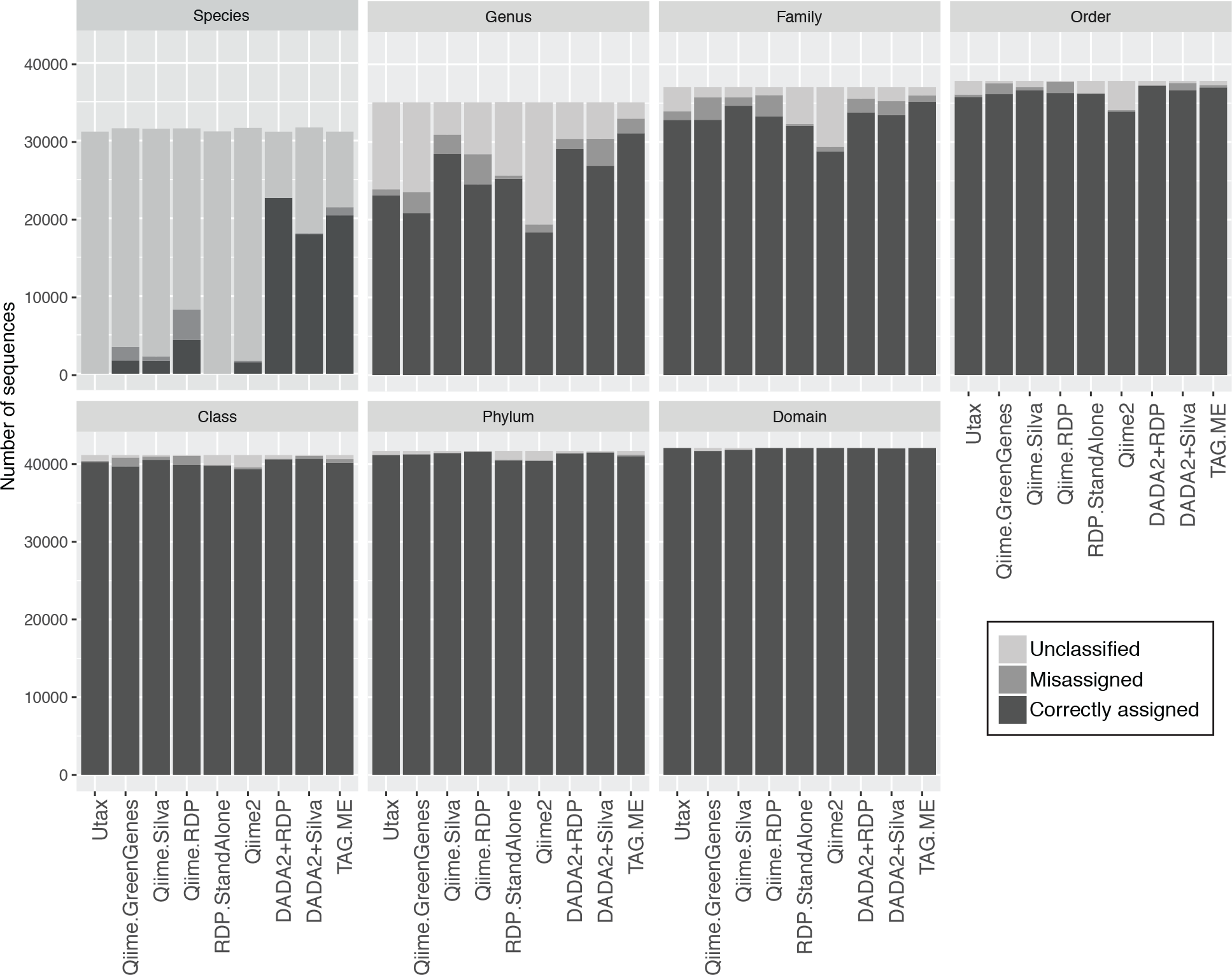
Comparison of TAG.ME taxonomic assignment with other methods.

The same dataset was used to evaluate the running time performance. While DADA2 took 1242 seconds to classify the complete dataset – 42069 sequences - up to species level, TAG.ME took 390 seconds in a 40-core workstation with 64GB of RAM. The proportion maintains as the number of sequences changes: DADA2 takes 403, 675, and 930 seconds while TAG.ME takes 100, 201, 298 seconds to classify 10000, 20000 and 30000 entries, respectively.

### Mock community

TAG.ME, DADA2 and QIIME2 Naïve Bayes were used to assign taxonomy to 124 different sequences representing the 515F-806R region of the V4 10 hypervariable region of the 16S rRNA gene from 102 different species. Based on this benchmark, TAG.ME presented a better performance, as represented in Figure 3A, correctly classifying 80 sequences, while DADA2 achieved 57 correct assignments, and QIIME2 only 10 classifications. The imprecise classifications are those that could not reach the deepest level: species. As QIIME2 reported a small number of classifications on species level, the method will not be discussed in this section, but the Supplementary Table 3 shows the assigned ranks for each sequence. Although the numbers are similar - 29 for TAG.ME and 28 for DADA2 - the depth of the imprecision is different since DADA2 have more classifications only until family and order levels, as shown in Figure 3B. In some occasions, both tools classified sequences from the mock community, but the reference species was unknown. The absence of reference made the comparison for these sequences impossible, and they were reported as “ref unknown” in Figure 3. One particularity must be addressed while discussing DADA2 species assignment: the possibility of multiple species classification. DADA2 uses exact sequence matches and this, sometimes, can assign more than one classification to one sequence. This fact can be interpreted as imprecision in the assignment or, without the required attention, can lead to misclassifications. The multiple species classification was observed in 30 sequences. If not considering the multiple species classification, both methods had few misassignments. The Figure 3C shows the dynamics of compared classification. Among the 80 sequences correctly classified by TAG.ME, 46 were also correct in DADA2, but 19 were imprecise, and 15 had multiple species assigned. DADA2 could correctly classify seven sequences that TAG.ME could not resolve, but 14 had multiple species that could lead to misclassification at the species level. We observed three sequences that could not be classified to deep levels because the reference was not included in the TAG.ME model. The Figure 3C also reinforces the precision and wariness of TAG.ME assignments. The complete comparison is available in Supplementary Table 3.

**Figure 3.**
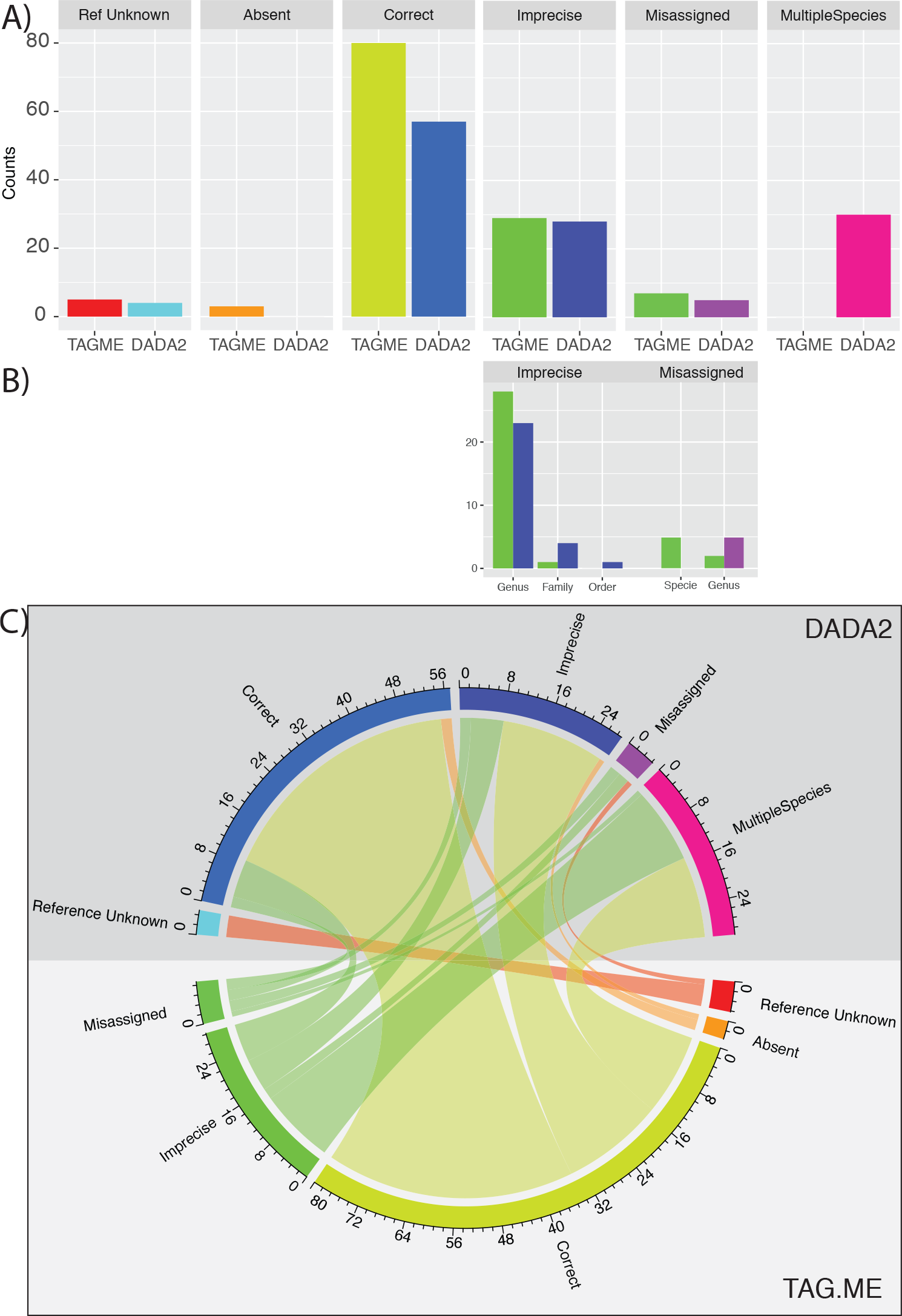
The comparison of the TAG.ME and DADA2 taxonomic assignments for the Mock Community benchmark dataset. (A) Barplots represent the number of sequences in each type of classification: “Ref unknown” refers to classified sequences that the mock community does not classify the species; “Absent” consists taxa that are absent in TAG.ME model construction; “Correct” comprises the taxonomic assignments that match the mock classification; “Imprecise” is the classification that could not reach the species level;“Misassigned” describes the assignments different from the mock reference; and “Multiple Assignments” are reads with more than one species assigned by DADA2. (B) The imprecisions and misclassifications were compared according to the taxonomic level of the last assignment. (C) The chord plot displays the inter-relationship between the sequence assignments.

### Gut and root microbiomes

TAG.ME performance was further compared with DADA2 on two real-world scenarios involving gut [21] and roots samples [22]. In these cases, both DADA2+RDP and DADA2+SILVA versions were compared. Figure 4A depicts the percentage of assignments per taxonomic level for gut samples, while Figure 4B shows the results for the root samples. TAG.ME capabilities become increasingly evident as more specific taxonomic levels were inferred. For gut microbiome samples, TAG.ME assigned almost twice as many sequences in comparison with DADA2 on Species level. The performance difference for the root sample was even higher. We observed that taxonomic assignment for ranks above species was slightly better when using DADA2+SILVA. The difference observed in this database is explained by the high abundance of classifications to taxa that still need more description and information (e.g., 0319-6G20, Elev-16S-1332, BIrii41, ABS-19, cvE6 - for Family rank, among others). TAG.ME is trained with a filtered database that removes these taxa that are not informative until the genus level - which is the case of the mentioned families - or have few entries to train the prediction model, and these entries were not present in the TAG.ME assignment. Although the percentage of assigned reads is not a measure of the classification quality, it is important to observe the differences in the proportion of assignments. Considering that, in the benchmark experiment, DADA2 and TAG.ME displayed similar accuracy, we would expect this test with biological samples reflecting that observation.

**Figure 4.**
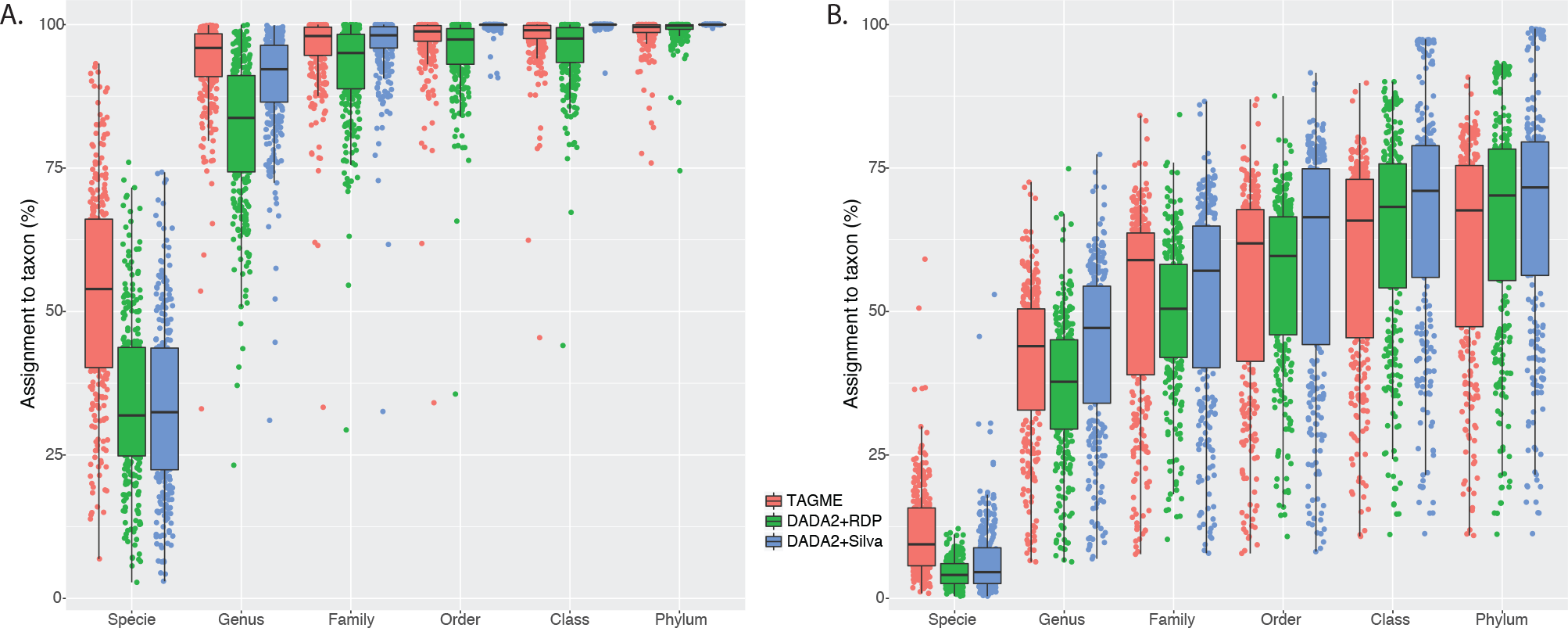
Comparative performance of TAG.ME and DADA2 on human gut (A) and root samples (B).

The rise of single variants - e.g., ASVs [8], ZOTUs [24] - supports the method used by DADA2, which is based on exact sequence match. DADA2 will have great performance, especially for well-described environments, with the engagement to produce more reference genomes. This is a fact observed when human microbiota is sequenced since the Human Microbiome Project encouraged the production of thousands of new reference genomes. Despite the efforts to build larger reference databases, we still can not picture many poorly explored environments, identify and represent all the microorganisms in one biofilm, or keep pace with the evolution process that will keep generating more and more variants. An alignment-free method, such as TAG.ME, with similar sensitivity and specificity as DADA2, is a relevant alternative to decide if one new ASV fits into one specific taxa pattern regardless of some small variation in the sequence.

### Limitations

Since TAG.ME relies amplicon specific models built using the tetranucleotides frequencies; it is recommended that the input sequences correspond to the whole amplicon generated by the pair of primers used to train the models. The use of partially sequenced amplicons decreases the frequency of some tetranucleotides and could lead to the reduced efficacy of our method. In this sense, it is highly suggested that the data shall be provided by Illumina sequencing, since it has fixed number of cycles reflecting on exact sequence length. The usage of this sequencing platform is a trend, and it is also suggested for DADA2 analysis. This is another point that makes TAG.ME complementary to the DADA2 workflow.

## 6. Conclusions

We show TAG.ME is an efficient and effective method for taxonomy assignment, capable of classifying well until species level, displaying good performance when compared to well-established methods. TAG.ME can be executed in a low-spec laptop with as little as 8GB of RAM memory, outperforming DADA2 in both run time and memory usage.

TAG.ME uses a multilevel supervised learning approach to assign taxonomy achieving not only a better assignment performance but also using considerably less computational resources. We believe our method will enable researchers to assess bacterial community structure and composition better, leading to novel insights regarding their role in nature and how to modulate these communities to our benefit.

TAG.ME is available as an open-source R package (https://gabrielrfernandes.github.io/tagme, Supplementary Software) that accompanies complementary analysis workflows and tutorial making TAG.ME fully integrated with other well-established pipelines, such DADA2’s.

## 7. Availability and requirements

**Project name**:TAG.ME

**Project home page**: http://gabrielrfernandes.github.io/tagme/

**Operating system(s)**: Multiplatform

**Programming language**:TAG.ME is an R package. The standalone scripts to build the models locally are written in Perl and R.

**Other requirements**: R

**License**:GNU General Public License

**Restriction**:None

***Availability of data and materials:*** The scripts used to build the models, and commands to execute them, are available for download at:

***Scripts and Tutorials***: https://gabrielrfernandes.github.io/tagme/

## 8. Abbreviations

OTU: Operational Taxonomic Unit
ASV: Amplicon Sequence Variant
AUROC: Area Under the Receiver Operator Characteristic Curve
LCA: Last Common Ancestor
ZOTU: Zero-radius Operational Taxonomic Unit

## 9. Author Contributions

D.E.V.P designed the experiment, supervised the research and wrote the paper. F.S.O and F.B.C implemented de algorithm, performed the analysis. D.K.M performed the analysis. G.R.F designed the experiment, implemented de algorithm, performed the analysis, wrote the paper.

## 10. Acknowledgements

We thank the Bioinformatics Platform of the René Rachou Institute − Fiocruz Minas − for providing the computational resources to develop and host this project. We also acknowledge the FAPEMIG support for FSO during the stay at the University of Pennsylvania.

## 11.

Author Information

1 Instituto René Rachou, Fiocruz Minas, Av. Augusto de Lima, 1715, 30190-009, Belo Horizonte, Brazil. 2 Post-Grad Program in Bioinformatics, Universidade Federal de Minas Gerais, Av. Antônio Carlos, 6627, 31270-901, Belo Horizonte, Brazil.

## Supplementary Material

**Supplementary Table 1.**
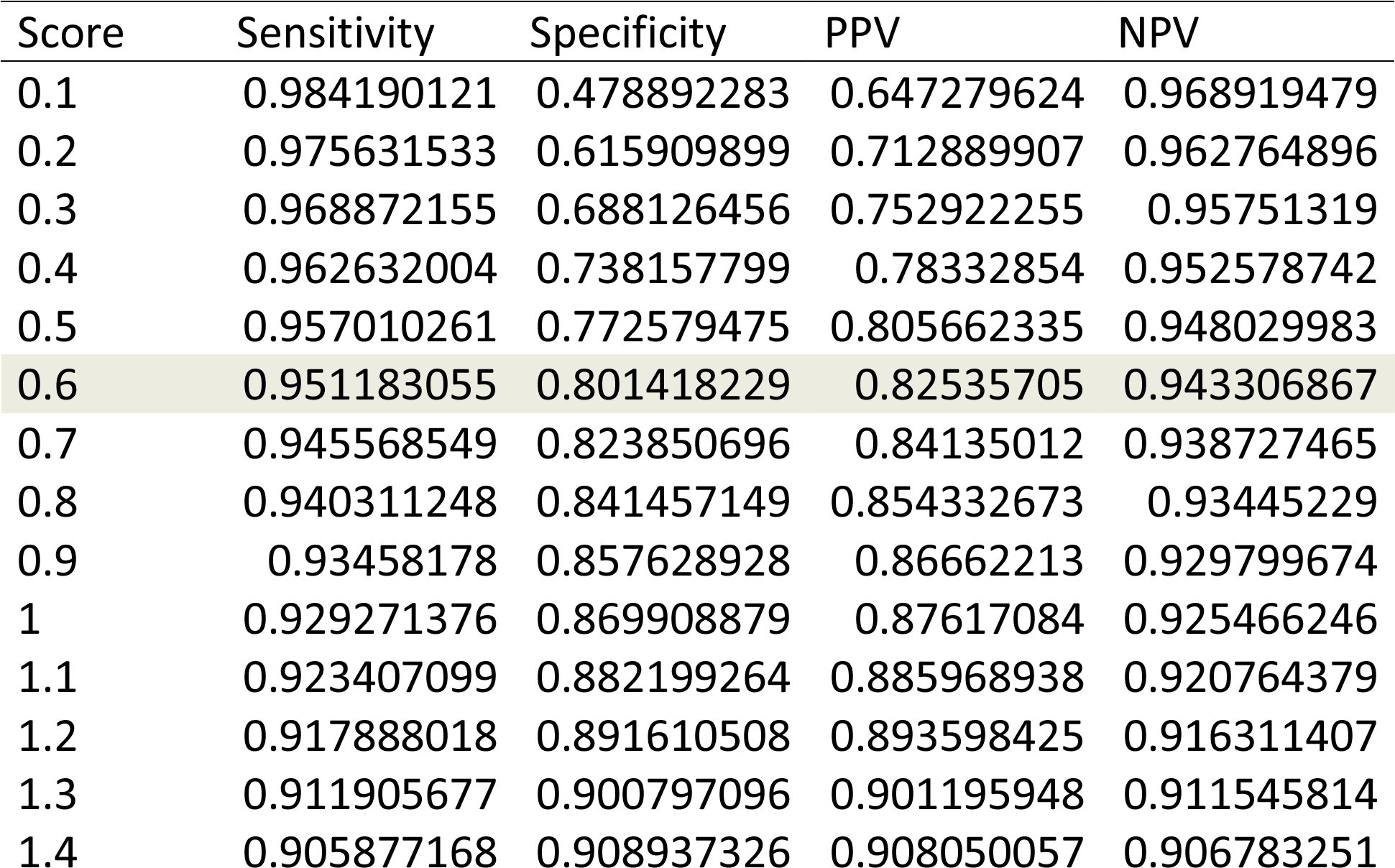
Specificity and Sensitivity of the genus model for the amplicon 515F-806R according to the Score. The highlighted row represents the specificity of 80%, resulting in a score threshold of 0.6 for classification.

Supplementary Table 2: https://figshare.com/s/ad950a745ef4aa846505.

Supplementary Table 3: https://figshare.com/s/a2eb466c808867e6f9f8.

**Supplementary Figure 1.**
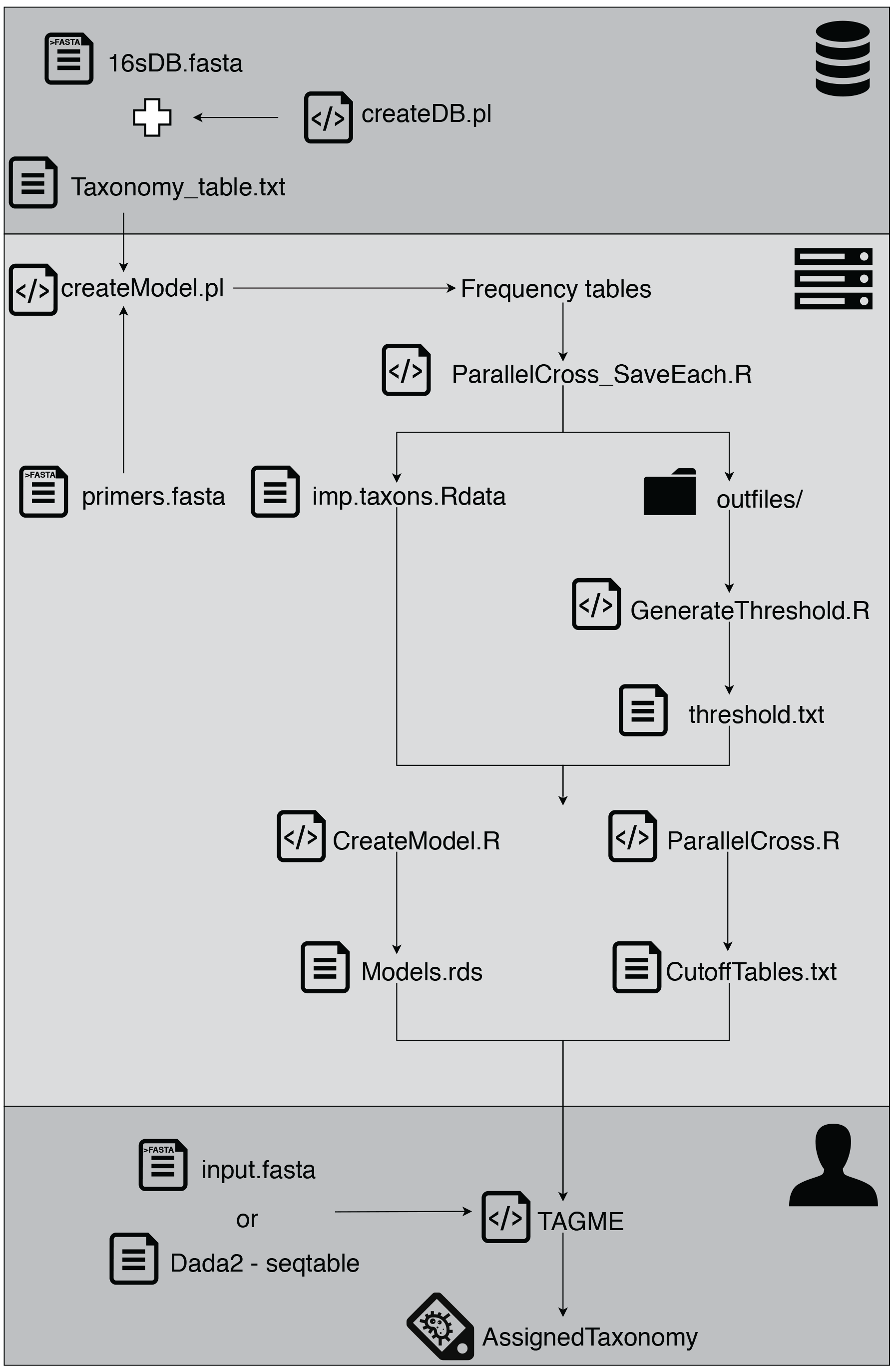
TAG.ME workflow to create the predictive models

**Supplementary Figure 2.**
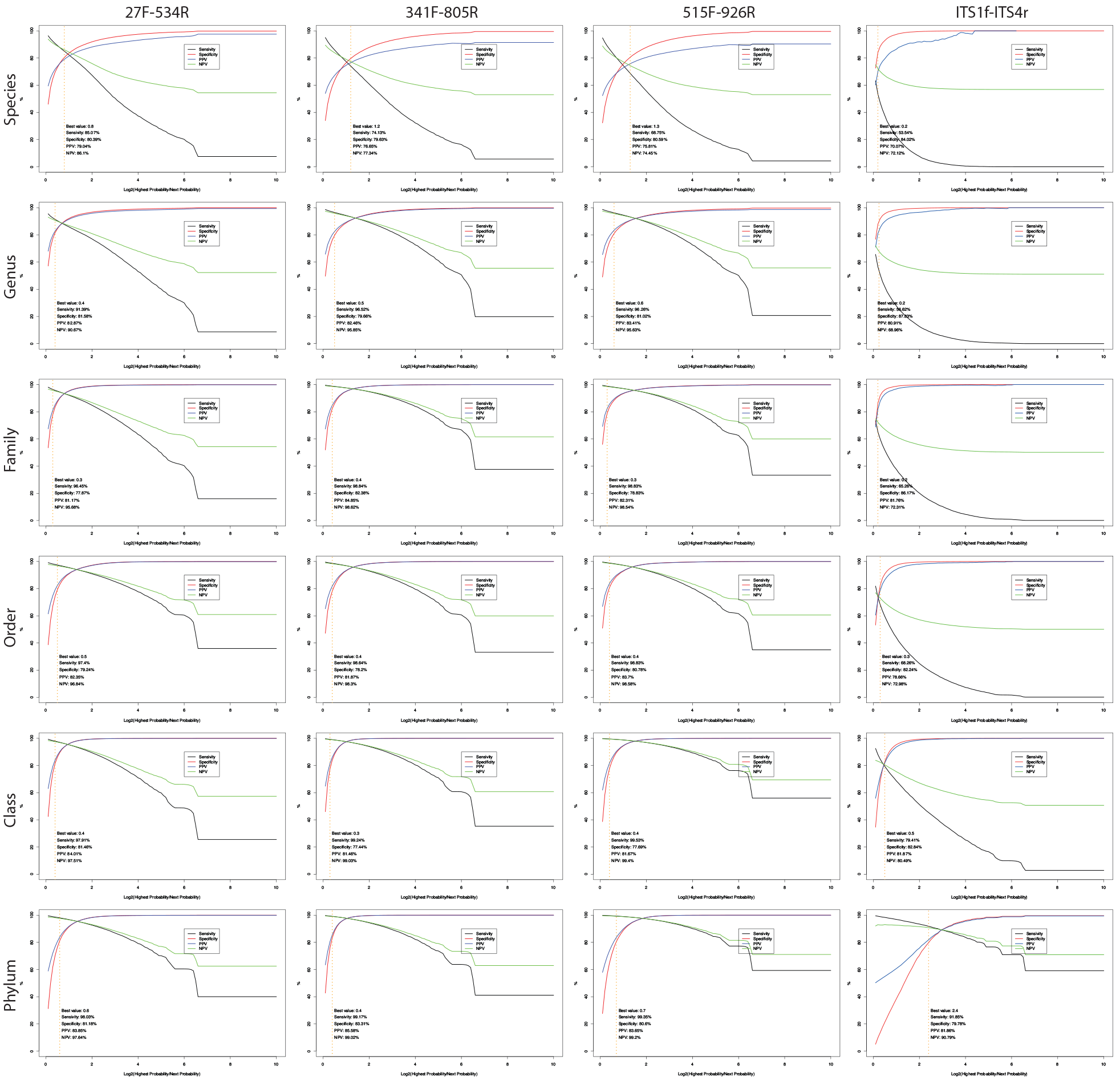
Performance measures – sensitivity, specificity, positive predictive value and negative predictive value – of each taxonomic rank for 2 different marker genes: 16s rRNA – 27F-534R, 341F-805R, and 515F-926R representing regions V1-V3, V3V4 and V4V5, respectively; and ITS.

**Supplementary Figure 3.**
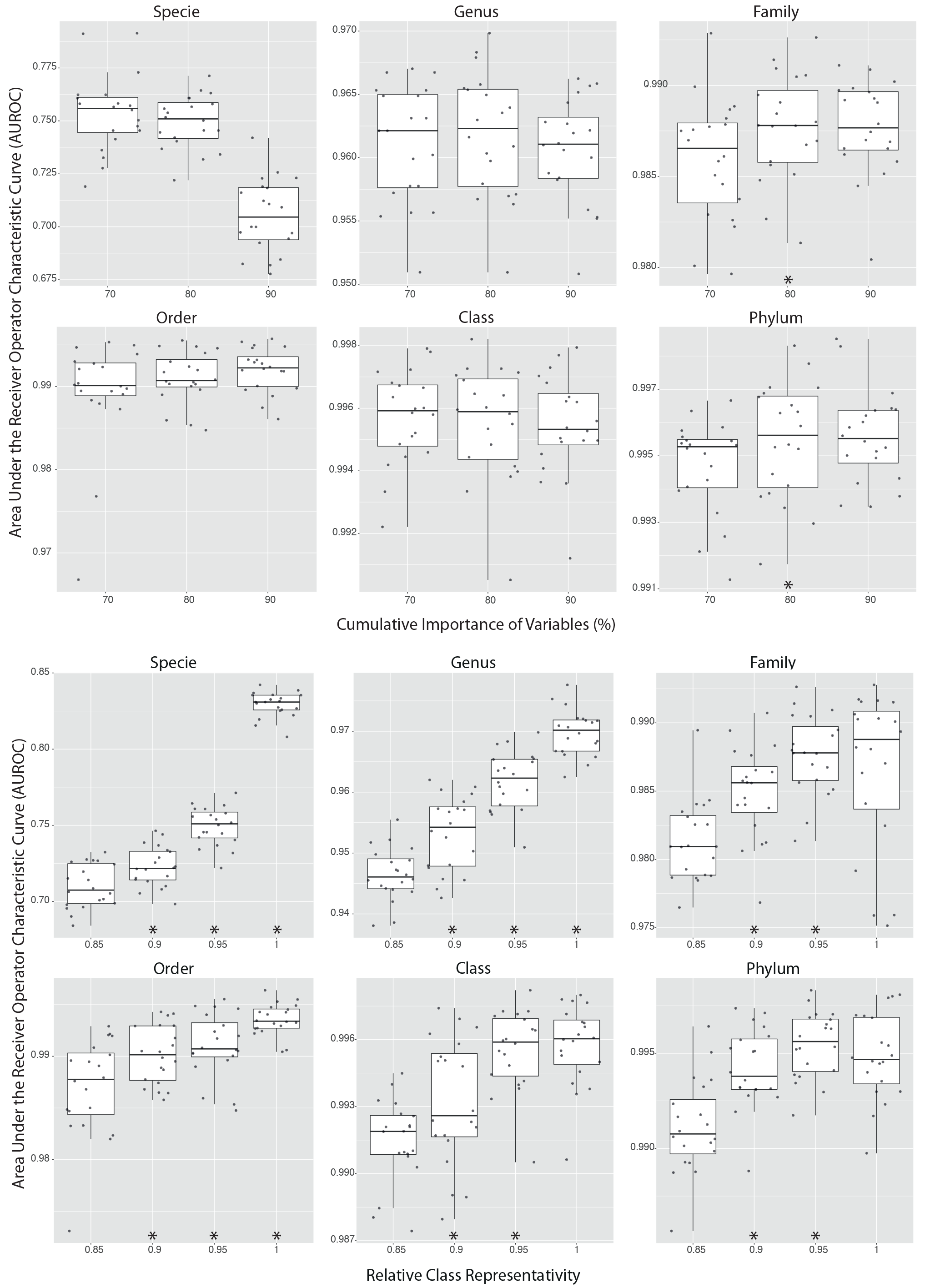
Distribution of AUROC across different tested parameters of (A) cumulative importance of variables and (B) relative class representativity. The * indicates significant (p<0.05) differences to support the bigger parameter selection.

**Supplementary Figure 4.**
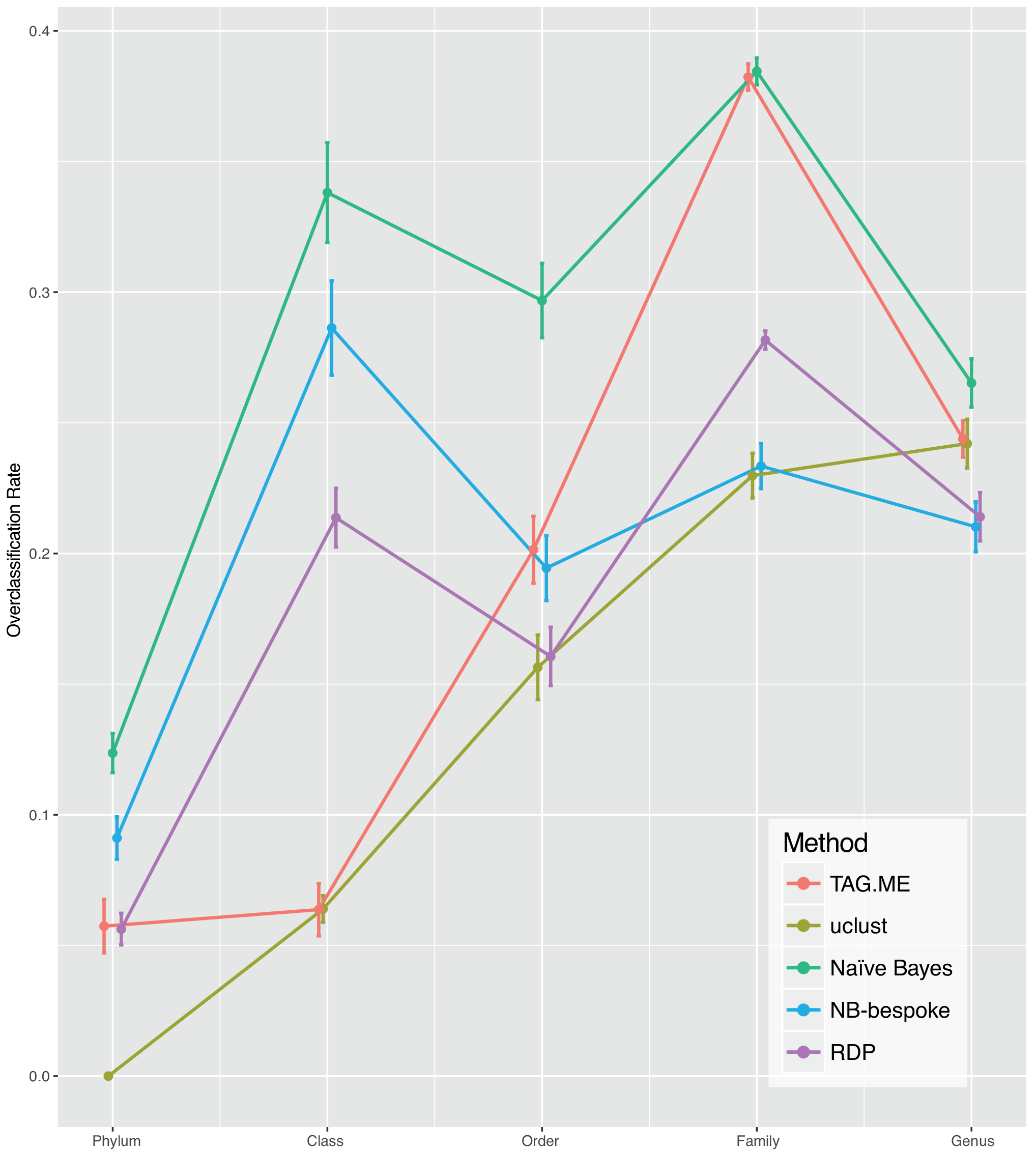
Comparison of the overclassification rate for each taxonomic rank among TAG.ME and QIIME2 methods: UCLUST, Naïve-Bayes, Naïve-Bayes bespoke and RPD.

